# ImputeCoVNet: 2D ResNet Autoencoder for Imputation of SARS-CoV-2 Sequences

**DOI:** 10.1101/2021.08.13.456305

**Authors:** Ahmad Pesaranghader, Justin Pelletier, Jean-Christophe Grenier, Raphaёl Poujol, Julie Hussin

## Abstract

We describe a new deep learning approach for the imputation of SARS-CoV-2 variants. Our model, ImputeCoVNet, consists of a 2D ResNet Autoencoder that aims at imputing missing genetic variants in SARS-CoV-2 sequences in an efficient manner. We show that ImputeCoVNet leads to accurate results at minor allele frequencies as low as 0.0001. When compared with an approach based on Hamming distance, ImputeCoVNet achieved comparable results with significantly less computation time. We also present the provision of geographical metadata (e.g., exposed country) to decoder increases the imputation accuracy. Additionally, by visualizing the embedding results of SARS-CoV-2 variants, we show that the trained encoder of ImputeCoVNet, or the embedded results from it, recapitulates viral clade’s information, which means it could be used for predictive tasks using virus sequence analysis.

## 1 Introduction

Since the start of the COVID-19 outbreak and the identification of the Severe Acute Respiratory Syndrome coronavirus 2 (SARS-CoV-2), the virus causing the COVID-19 pandemics, laboratories around the world have been generating viral genome sequence data with unprecedented speed, enabling real-time progress in the understanding of this new disease and in the research and development of candidate medical countermeasures [1, 2]. This step is necessarily needed because sequence data are essential to design and evaluate diagnostic tests, to track and trace the ongoing outbreak, and to identify potential intervention options. Nevertheless, following the rapid explosion of SARS-CoV-2 sequences, imputation of detected genetic variants worldwide is becoming a major priority, as the number of missing nucleotides spread across the available data is quite large and hampers a comprehensive study of the available samples. Yet, reliable imputation of missing nucleotides in genomic sequences has always been a challenging problem in biology owing to the fact that genomic sequencing is an imperfect process. This study aims at the efficient and reliable imputation of such samples with the help of recent advances in deep learning [3–5].

## 2 Previous Studies

In general, the imputation methods can be divided into two major categories: (1) reference-based, and (2) reference-free. If an imputation method is based on sequences and what is found within the given populations in order to correctly assign a missing nucleotide, then the method is a reference-based method of imputation. Distance-based methods (e.g., using Levenshtein distance [6] or Hamming distance [7] to identify the closest sequence to an incomplete sequence in a reference dataset) are examples of such type of imputation. They work very well in sequences that do not undergo recombination, but they are quite slow to run. On the other hand, if the method is focused on pattern recognition, then the imputation method would be considered to be reference-free. ImputeCoVNet, proposed in this study, is a deep learning reference-free approach for the imputation of SARS-CoV-2 sequences.

We distinguish our proposed deep learning network and the problem we address here from earlier studies which in particular worked on genotype imputation and mainly focused on human data [8–10]. There are also more recent approaches that employed deep learning techniques for human genotype imputation which are discussed in [11, 12].

However, concerned with SARS-CoV-2 sequences in our analysis for which imputed sequences are needed, and in contrast with human genotype data, it should be considered that the evolution of RNA viruses must be assessed from the perspective of the quasispecies defined as the population of genomes linked through mutations with each other. The initial genome sequence was identify in Wuhan in December 2019, and all newer sequences are generally aligned to detect variants. Since the initial sequencing of the SARS-CoV-2 genome, there has been over 10,000 mutated sites identified to be segregating within the human population, resulting in a mutant spectrum that interacts at the functional level and works as a selection unit [13]. Structurally, each individual genome in the spectrum can be defined as a collection of the alleles at the mutated sites, either showing the initial allele or a variant allele occurring on the same viral genome, which is known as a ‘haplotype’.

## 3 Genomic Data

### SARS-CoV-2 Data Source

The SARS-CoV-2 genomic data used in this study was submitted as part of the Global Initiative on Sharing All Influenza Data (GISAID) database^2^ [14]. GISAID is a reference database as its data submitters and curators ensure real-time data sharing of human coronavirus 2019 (hCoV-19) remains reliable for rapid progress in the understanding of the new COVID-19 disease. In addition to the SARS-CoV-2 sequences themselves, the samples are provided with their associated metadata which includes age, sex, collection date, submission date, exposed continent, exposed country and viral clade. We obtained the 35,698 high-quality consensus sequences available on July 22th 2020, and each sequence was aligned to the reference genome (NC_045512.2) using minimap2 v.2.17. Single nucleotide variants (SNVs) were called using samtools [15] and bcftools [16]. Low-quality regions were identified and mutations in low-quality regions were masked. We identified a total of 6,349 sites that were found in more than one sequence.

### Evaluation Data Preparation

As there existed no predefined test data (with ground truth) for the evaluation of SARS-CoV-2 data imputation models, in the first step we prepared such evaluation data. This data preparation consisted of a ‘missing position injection scheme’ applied to ‘complete’ haplotypes, which did not include any missing values, hence for which the ground truths were known (assuming no errors). This mandatory step resulted in three mutually- exclusive splits of data, namely train, validation, and test sets (80%-10%-10%) on which the neural network and distance-based (baseline) approaches were trained and evaluated. To ensure the test and validation data were as similar as possible to real ‘non-complete’ haplotypes and avoid any bias towards less problematic sites, the missing position injection scheme learned the missing position distribution of the non-complete haplotypes in advance. Due to stochasticity in the selection of haplotypes and missing sites, the data preparation step was executed 5 times and the mean was reported as the final accuracy results. This whole process was done for three different haplotype definitions, made of positions for which the variant allele frequencies were of 0.01 (49 sites), 0.001 (244 sites), and 0.0001 (885 sites). Figure 1 represents the distribution of missing position in the real data v.s. one of the data generation steps.

**Figure 1:**
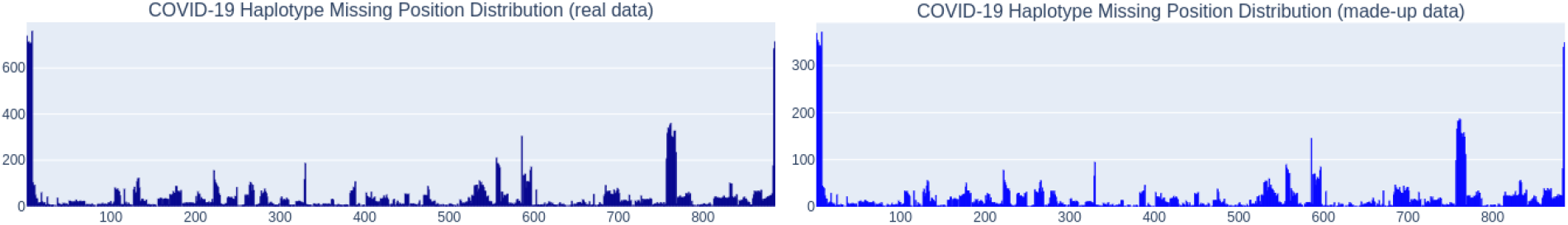
Comparison of the distribution of missing positions in real v.s. made-up data employed for evaluation of the imputation models (allele frequency of 0.0001) Additionally, for our second experiment, to prepare another set of test data to evaluate how imputation models perform for the imputation of the future test sequences when provided only with training sequences of an earlier time-point (May 1st, 2020), the train and validation splits were built from those specific predated haplotypes. For these data splits, the missing position injection scheme learned only from the predated non-complete sequences as well. For the test data, however, the injection process was executed on haplotype sequences collected after that specified time.

## 4 ImputeCoVNet Architecture

### Network Definition

While deep neural convolutional operations are mainly used in computer vision [17–19], an autoencoder is an unsupervised learning technique for neural networks that learns efficient data representations (encoding) of input data [20]. ImputeCoVNet is a 2D convolutional ResNet autoencoder that aims at learning and reconstructing the input SARS-CoV-2 haplotypes. The whole network consists of two sub-networks: (1) an encoder which is responsible to encode the given input into a low-dimensional vector determined in the bottleneck, and (2) a decoder that is responsible to reconstruct that sequence from that low-dimensional vector. Figure 2 illustrates the architecture of the ImputeCoVNet designed to address the imputation of SARS-CoV-2 sequences.

**Figure 2:**
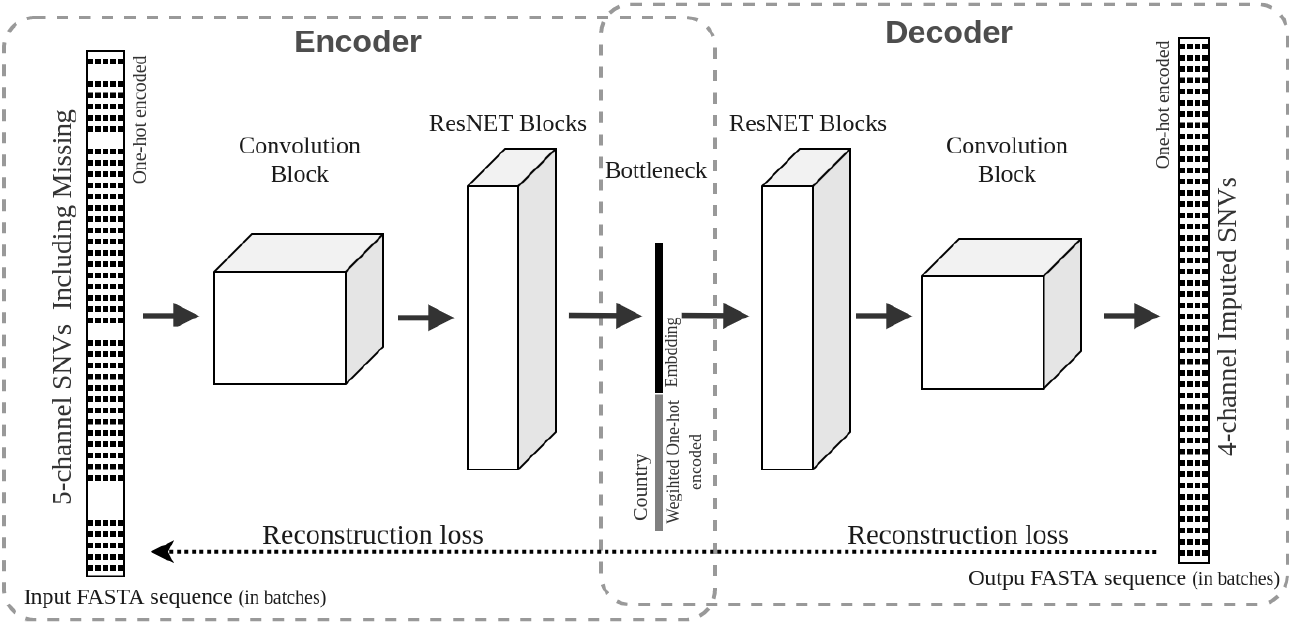
ImputeCoVNet architecture including a 2D ResNet encoder and a 2D ResNet decoder.

### Model Training

During network training, the first and last layer receive the complete input and output haplotypes as one-hot encoded representations of their individual variants while the reconstruction loss aims at optimization of the network’s weights and biases. However, for proper hyper-parameter selections of the size of kernels, number ResNet blocks [21], learning rate, etc. the validation split (non-complete haplotypes) was employed and imputed. The imputation accuracy of the validation split determined the final hyper-parameters of the network. For the final evaluation on the test split, we combined train and validation split together (complete haplotypes) and trained the model on them from scratch, and used that model to impute the test haplotypes. As an implementation preference, since at test/validation time there would be missing values, we considered an extra channel for them in the one-hot encoded representation used by the network; however, as an alternative, those missing positions could be represented as all-zeros (in contrast to one-hot encoded) in their encoded vectors.

Additionally, we investigated if the provision of metadata to decoder, along with the encoded variant, improved the imputation accuracy of missing sites (applied during model training and testing). For this purpose, the metadata vector that is typically provided as a (weighted) one-hot encoded representation of the metadata of interest (e.g., exposed country), was concatenated to the embedded sequence resulting from the encoder before passing them to the decoder as shown in Figure 2.

### 5 Evaluation and Results

In our evaluation of ImputeCoVNet, we considered an approach based on Hamming distance (HD) as the baseline model. To compute the results from the HD approach, the combination of train and validation splits were considered as the reference data (their complete haplotypes) to which the distances of non-complete haplotypes (query) in the test split were calculated. For each query, the reference haplotype with the smallest distance was selected for the imputation and the missing sites in the query were replaced by the selected haplotype’s alleles. Whenever there was a tie in distances to two winning reference haplotypes (with the least distance to the query) the one with higher occurrence in the reference dataset was selected for the imputation of the missing sites in the query.

### Evaluation on current non-complete sequences

The first evaluation experiment was conducted for the imputation of haplotypes where the training data (or reference data) and the test data were from the same time-span (ie. from December 2019 to July 2020). Table 1 presents the results of this experiment.

**Table 1:**
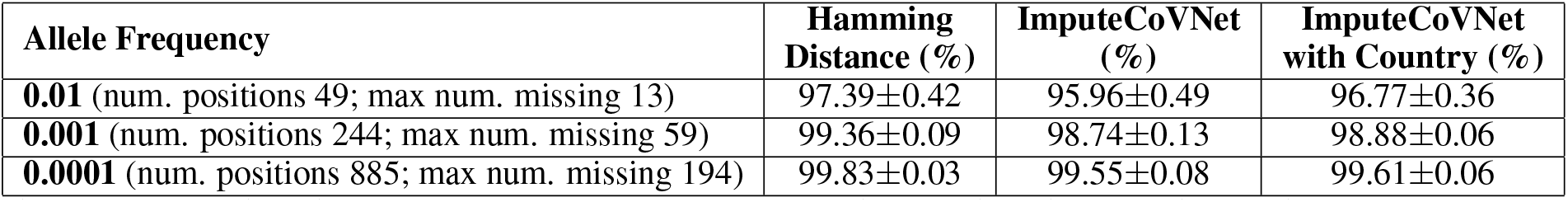
Position-based imputation accuracy comparison of distance-based approach v.s. deep learning network ImputeCoVNet, when evaluated on SARS-CoV-2 haplotypes composed of variants with three different allele frequency thresholds.

The results from ImputeCoVNet are comparable to the computationally expensive imputation resulting from Hamming distance, with both approaches reaching high accuracy, which improves with the number of variants considered. This can be due to improvement of imputation because of longer SARS-CoV-2 haplotypes, or can reflect the fact that the likelihood of detecting an error at sites with low-frequency variants is lower. Since ImputeCoVNet is more sensitive to the number of training examples from the pattern recognition perspective, we expect that, as the number of SARS-CoV-2 sequences samples increase, the results from ImputeCoVNet would be on par with or better than the HD algorithms. Notably, for the imputation of the test split, the average imputation time for haplotypes with an allele frequency of at least 0.0001 (885 variants) was only 6 seconds for ImputeCoVNet, whereas the HD approach took 1 hour and 25 minutes to impute the same split. Also, we observed in all cases that the provision of metadata to the decoder improves the prediction of missing alleles. This improvement is more considerable in predicting variant with allele frequency over 0.01 (49 variants). In our experiments, we noticed the ‘exposed country’ improved these results more than ‘exposed continent’ while the other metadata considerations such as sex, age and date had no measurable impact at this stage of the pandemics.

### Evaluation on future non-complete sequences

We restricted our second experiment to the model trained to impute variants with allele frequency above 0.0001. Table 2 shows the results for this experiment in which the training data is from the past and the models are tested on future sequences.

**Table 2:**
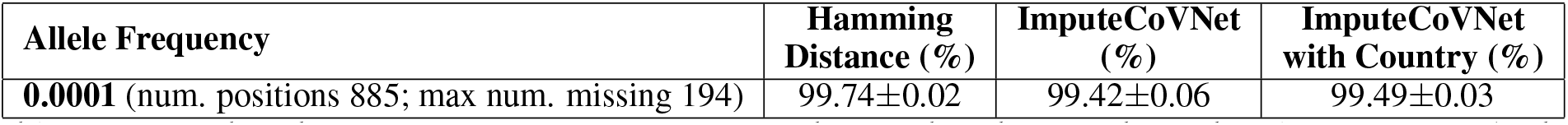
Position-based imputation accuracy comparison of distance-based approach v.s. deep learning network when evaluated on allele frequency of 0.0001 of future SARS-CoV-2 sequences.

| Allele Frequency | Hamming Distance (%) | ImputeCoVNet (%) | ImputeCoVNet with Country (%) |
| --- | --- | --- | --- |
| **0.0001** (num. positions 885; max num. missing 194) | $99.74 \pm 0.02$ | $99.42 \pm 0.06$ | $99.49 \pm 0.03$ |

We did not notice any significant drop in the imputation results, which means that ImputeCoVNet approach is expected to have the same accuracy for imputing sequences generated in the next months. However, this will have to be reevaluated as sequences from the second wave pour in. Compared to the previous experiment, the slight decrease in the accuracies could be also explained by fewer training samples in this experiment due to the chosen time-point (25,630 sequence samples here v.s. 26,663 in the previous experiment).

### PCA representation of embedded sequences

Due to the naturally expanding genetic diversity of SARS-CoV-2 viruses, GISAID introduced a nomenclature system for major viral clades, based on marker mutations within 6 high-level phylogenetic groupings from the early split of S and L, to the further evolution of L into V and G and later of G into GH and GR^3^. Figure 3 represents the principal component analysis (PCA) of the embedded SARS-CoV-2 test haplotypes in the bottleneck once they were passed through the encoder network, colored with respect to these GISAID clades. These embeddings exclude the country vector and were computed when ImputeCoVNet were fully trained.

**Figure 3:**
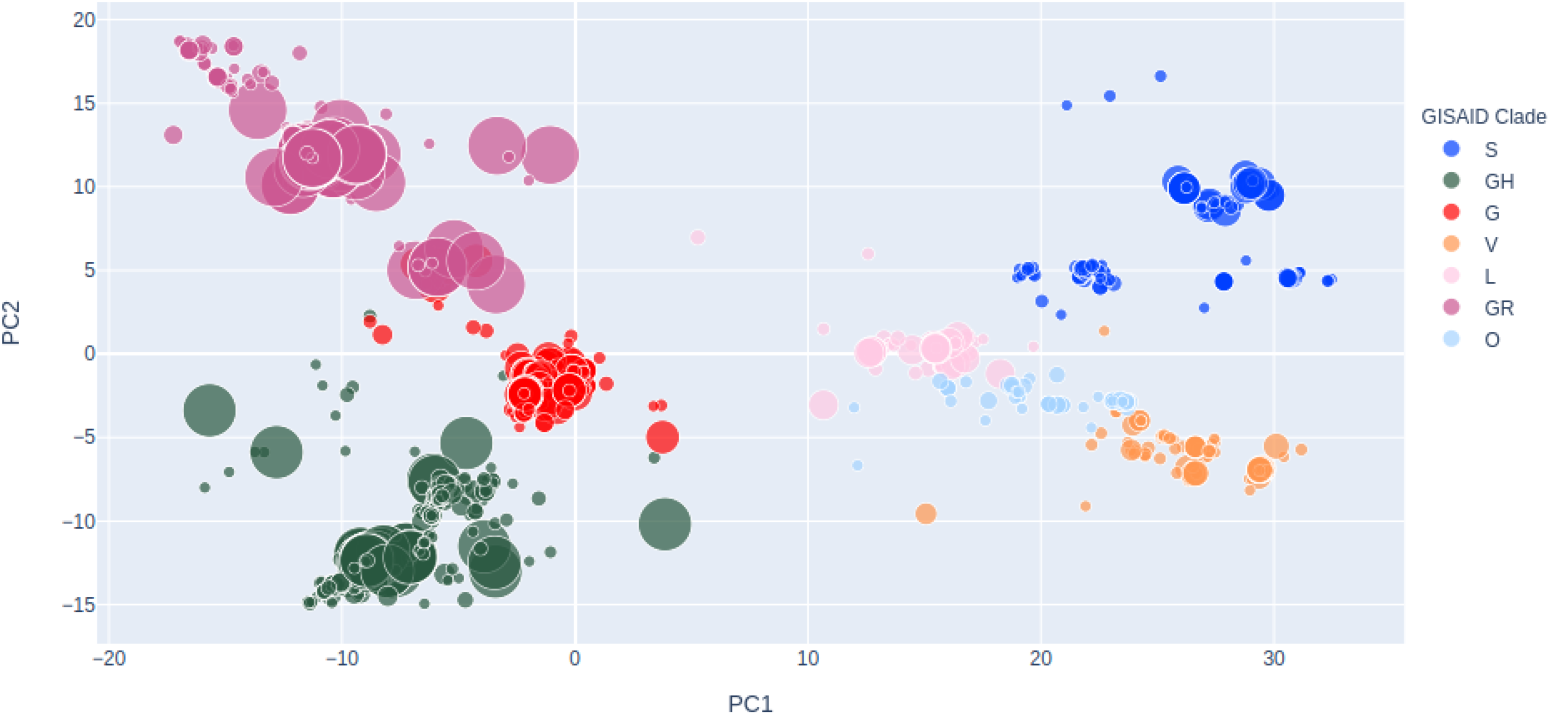
PCA representation of the SARS-CoV-2 sequences embedding vectors resulted from the the encoder network

Figure 3 shows the ImputeCoVNet is able to extract highly relevant features of given SARS-CoV-2 sequences with the help of an encoder and through the non-linearity functions. This feature extraction scheme suggests that ImputeCoVNet can be employed for predictive tasks in virus sequence analysis while including metadata as well, particularly when the number of labelled data is limited.

## 6 Conclusion

This paper proposed an autoencoder deep learning network architecture named ImputeCoVNet that aimed at fast and reliable imputation of SARS-CoV-2 variants. When compared with the Hamming distance approach, the imputation accuracy results were comparable and very high, especially when very low-frequency alleles were included. We also presented that the provision of metadata to the decoder network improves imputation results. While the distance-based approach tends to demand more processing time for imputation as the number of haplotypes to impute increases, deep learning models such as ImputeCoVNet, once trained, execute imputations almost in real-time. No clear drop in the accuracy has been seen when testing on data generated exclusively after the training sequences, this algorithm could therefore be deployed to impute all the new sequences submitted to GISAID.

## Footnotes

2 https://www.gisaid.org

3 https://www.gisaid.org/references/statements-clarifications/clade-and-lineage-nomenclature-aids-in-genomic-epidemiology-of-active-hcov-19-viruses/

